# An efficient biochemical method for characterizing and classifying potentially amyloidogenic and therapeutic peptides

**DOI:** 10.1101/2024.05.06.592687

**Authors:** Lucas Pradeau-Phélut, Stacy Alvès, Léo Le Tareau, Cyann Larralde, Emma Bernard, Josephine Lai Kee Him, Eléonore Lepvrier, Patrick Bron, Christian Delamarche, Cyrille Garnier

## Abstract

Amyloidosis are proteinopathies characterized by systemic or organ-specific deposition of proteins in the form of amyloid fibers. Nearly forty proteins have been identified to play a role in these pathologies and the structures of the associated fibers are beginning to be determined by Cryo-EM. However, the molecular events underlying the process, such as fiber nucleation and elongation, are poorly understood, which impairs developing efficient therapies. In most cases, only a few dozen amino acids of the pathological protein are found in the final structure of the fibers, while amyloid peptides comprising 5 to 10 amino acids are involved in fibers nucleation process. The identification and biochemical characterization of these peptides is therefore of major scientific and clinical importance. *In silico* approaches are limited due to the peptides’ small size and long-distance intra- and intermolecular interactions that occur during nucleation. To address this problem, we developed a novel biochemical method for characterizing and classifying batches of related peptides. Initial work to optimize our approach is based on the reference peptide PHF6 (ß1) from Microtubule Associated Protein Tau (MAPT) as compared to 22 related peptides. We classified these peptides into groups displaying different biochemical properties, and thereby identified new amyloid peptides and peptides with therapeutic potential. We underline that our method is applicable to any family of peptides and could be scaled up for high-throughput analyses.

## Introduction

Amyloid pathologies, also called amyloidosis, are characterized by the abnormal deposits in tissues. Since these deposits display a blue color in the presence of iodine common with starch, they are commonly referred to as “amyloid” (Virchow, 1856). It has been demonstrated that amyloid deposits possess several characteristics. For example, they bind Congo Red enabling them to emit green birefringence when subjected to polarized light (Bennhold, 1922). Likewise, the fluorescence of Thioflavin T is exacerbated when it intercalates into amyloid deposits (Biancalana and Koide, 2010; LeVine, 1993; Naiki et al., 1990). Thioflavin T was first described in 1959 (Cohen and Calkins, 1959) and is now the reference dye for amyloid structures due to its high sensitivity (Biancalana and Koide, 2010; Naiki et al., 1990; Xue et al., 2017). In 1959 electron microscopic examination of ultrathin sections of amyloidotic tissues revealed the presence non-branching fibrillar structures 8 to 10 nm in width (Cohen and Calkins, 1959). The protein nature of these deposits was revealed later in the 1960s when it was demonstrated by X-ray diffraction that the deposits displayed an organized β-sheet structure, showing typical southern and equatorial diffraction spots of 4.7 Å and 11 Å (Glenner et al., 1969). Amyloid deposits can be made up proteolytic fragments of protein, for example in the case of Amyloid Precursor Protein (APP) or mutated Fibrinogen A α-chain, peptides of around forty amino acids lead to the formation of amyloid deposits (Borchelt et al., 1996; Garnier et al., 2017a). When fibers are made up of entire proteins, only a small fraction of them (comprising few tens amino acids) is present in the core of the fibers structures (Fitzpatrick et al., 2017). Approximately 40 proteins have been recognized as amyloidogenic by the ISA Nomenclature Committee, while others are in the process of being validated (Buxbaum et al., 2022). They include immunoglobulin light chains involved in L-amyloidosis, certain circulating blood proteins, *e*.*g*. Apolipoproteins E and SAA, associated with amyloidosis A, or transmembrane proteins such as APP or even soluble cellular proteins such as Tau involved in Alzheimer’s disease. These proteins have no real common features except for their singular amyloidogenic potential, which leads to self-association in amyloid structures. The causes underlying to the loss of protein function differs between pathologies. However, genetic factors, excessive protein concentration due to a dysregulation of its cellular homeostasis, absence of structures or misfolding, incomplete proteolysis or post-translational modifications have been suggested (Buxbaum and Linke, 2012). In most cases, several factors, often two, seem to be simultaneously responsible for triggering the pathology. Whatever the causes or the type of mechanism involved in fibrillation are, a common feature among these auto-association processes is the loss of function of the original protein. The protein aggregates cause more or less progressive and often severe pathologies, regardless of whether the deposits accumulate systemically in the organism or in a specific organ. Therefore, the identification of the proteins involved and a better understanding of their self-associative processes are essential for effective patient management and/or the discovery of inhibitors. For example, in a previous study, we discovered the involvement of fibrinogen Aα in a kidney amyloidosis which allows for adapting patient management. In addition, the identification of the amyloid pentapeptide “**<u>VLITL</u>**” responsible for the pathology will ultimately make it possible to identify an inhibitor (Garnier et al., 2017a). In most cases, the initial nucleation process involves short peptide sequences containing 5 to 10 amino acids. For example, within aß42 peptide, two sequences, **<u>IIGL</u>** and **<u>GVVI</u>**, contribute to its amyloidogenicity (Sgourakis et al., 2007). For the Tau protein, the hexapeptide _**<u>306</u>**_**<u>VQIVYK</u>**_**<u>311</u>**_ (ß1) seems to play an initiating role in nucleating fibers *in vivo* as well as *in vitro* (Arakhamia et al., 2021; Falcon et al., 2018; Fitzpatrick et al., 2017; Perez et al., 2007; Schweighauser et al., 2023; Zhang et al., 2019).

The study of these short highly amyloid peptides and their characterization and classification holds significant interest to understand their key role in the nucleation processes. This requires implementation of various complementary approaches, including predictive modeling and experimental techniques such as biochemical, biophysical and structural biology.

In this study, we optimized biochemical methods to characterize and classify a batch of peptides related to a known amyloid peptide. Depending on their biochemical properties the target peptides were located on a Garnier-Delamarche plot (GD-plot). The positioning of a given peptide depends on its aggregative character (x-axis, linear scale) and on its amyloid character compared to the reference peptide for which fluorescence was set to 100% (y-axis, logarithmic scale). We note that the approach can be automated and allows for classification of any batch of related peptides produced from a reference peptide. To improve the method, we employed Tau’s hexapeptide PHF6 (_306_VQIVYK_311_: ß1) as a reference. We selected 22 related peptides that have the same amino acid composition as ß1 (I, K, Q, 2V, Y) but different sequenses. The biochemical properties of the peptides and their positioning on the GD-plot enabled us to identify aggregative, amyloid and soluble (*i*.*e*. neither amyloid nor aggregative) cases. The biochemical properties of the peptides were compared to their predicted amyloid/aggregative features. Finally, we focused on peptides classified as amyloids, and we highlighted the interest of such characterization/classification may have for the development of future therapies.

## Materials & Methods

### Hexapeptides preparation and storage

The PHF6 (ß1) peptide and derivative hexapeptides were synthesized as Ac-[V;Q;I;V;Y;K]-NH2 form and lyophilized by Proteogenix®. The purity was verified using ESI-MS or MALDI-TOF MS. Peptide powder was resuspended in deionized water (Milli-Q, Merck) at a final concentration close to 5 mg.mL^-1^, vortexed and sonicated for 15 minutes each at 35°C in a sonication water bath (Advantage-Lab AL04-04) before being ultracentrifuged at 100,000 g for 15 minutes at 20°C. The pellets were removed and supernatant concentrations determined by measuring the UV absorption at 280 nm using a molar extinction coefficient of 1450 M^-1^.cm^-1^. Finally, the hexapeptide sample was aliquoted and stored at −80°C for further experiments (Schirmer et al., 2016).

### Prediction of hexapeptide amyloidogenic properties

The predictions were carried out using a virtual protein sequence containing concatenated hexapeptides separated from each other by a nonapeptide-spacer flanked between two glycines (**G**PNSKQSQDE**G**). This nonapeptide is part of DNA binding protein 43 (TDP-43 residues 178 to 186). This peptide was chosen as spacer because it was predicted to be non-aggregative (Garnier et al., 2017). We used Pasta 2.0 (Walsh et al., 2014), Tango (Fernandez-Escamilla et al., 2004; Linding et al., 2004; Rousseau et al., 2006) and Salsa (Zibaee et al., 2007) for predictions and normalized the values as described in (Garnier et al., 2017b). To compile results, scores for each amino acid were expressed as a percentage considering that the maximum score given by the method was 100%. Then, percentages obtained were averaged for each amino acid. The overall peptide score is an average of the scores for six amino acids. Finally, score values were normalized against PHF6 (ß1 reference) for which we set the score at 100%.

### Hexapeptide polymerization

After rapid thawing, hexapeptide aliquots were pre-diluted in deionized water and incubated for at least four hours at 25°C. Hexapeptides concentration was measured by UV absorption as indicated in the “Hexapeptide preparation” section, and adjusted to 1.25 times the working concentration. Hexapeptide polymerization was induced by adding of 5X fibrillation buffer (50 mM MOPS, 750 mM NaCl, pH = 7.2). Samples were polymerized for at least 1-2 hours at 25°C. Whenever polymerization processes were followed by spectrofluorimetry, the sample were supplemented with 32 μM Thioflavin T (Schirmer et al., 2016).

### Hexapeptides aggregative properties

Once polymerized, samples were ultracentrifuged at 100,000 g for 5 minutes at 25°C (Beckman TL100, rotor TL-A100). Supernatants were recovered and the hexapeptide concentrations of were determined by UV absorption as previously described. The concentration of sedimentable hexapeptide structures was deduced from the difference between supernatant and initial concentrations.

### Hexapeptides amyloidogenicity

Hexapeptides were polymerized at 800 μM. Once polymerization was finished, 40 μl of the sample was diluted in 360 μl of 1X fibrillation buffer containing 10 μM Thioflavin T in 0.2 x 1.0 cm thermostated black quartz cuvettes. Fluorescence spectra between 470 to 600 nm at an excitation wavelength of 450 nm (Perkin Elmer LS 55, PM 750, slits 10), were immediately acquired after the dilution and homogenization steps. For each experiment, a ß1 sample was used as control. Results were normalized considering that ß1 sample had a maximum fluorescence intensity of 100%. The maximum error measured on this type of experiment was +/-38%, it was applied to all experimental points.

### Amyloid hexapeptides, critical assembly concentrations and reversibility

To determine critical assembly concentrations (Cr), hexapeptides were polymerized at different concentrations for 1.5 hours as described above. After centrifugation, 16,900 g, 15 minutes at 25°C, supernatants were recovered and the concentration of ß1 was determined by UV absorption as described above. ß1 sedimentable concentration was deduced from the difference between supernatant and initial concentrations. The concentration of the sedimentable fraction was determined and plotted as a function of the total peptide concentrations. Crs were determined by using linear regression models and solving the equations y(x) = 0, where x corresponds to the Cr.

To assess reversibility, hexapeptides were polymerized at twice their Cr for 1.5 hours at 25°C. Reversion was induced by 8-fold dilution step with 1X fibrillation buffer 10 μM Thioflavin T to achieve a final concentration 4-fold lower than their proper Cr. Kinetics of depolymerization were monitored over time using fluorescence in 0.2 x 1.0 cm thermostated quartz cuvettes at emission wavelength of 485 nm with an excitation wavelength of 450 nm (Perkin Elmer LS 55, PM 750, the slits were adjusted depending on hexapeptide).

### Samples preparation for electron microscopy

Four microliters of hexapeptide samples were applied to carbon-coated copper grids (300 mesh), and allowed to stand for two minutes. The grids were then washed with 1X fibrillation buffer, negatively stained for 1 minute with 1% (w/v) uranyl acetate, and wicked dry prior to analysis. Observations were performed using a JEOL 2200 FS electron microscope operating at 200 kV and equipped with a 4k × 4k CCD camera. Images were captured at different magnifications (6,300X, 11,000X, 22,000X) and processed using ImageJ software.

For cryo-EM, 3 μL of hexapeptide sample were applied to glow-discharged Quantifoil R 2/2 grids (Quantifoil Micro tools GmbH, Germany), blotted for 1 second, and then flash frozen in liquid ethane using the semi-automated plunge freezing device CP3 (Gatan inc.) at 95% relative humidity. Images were acquired using a JEOL 2200 FS electron microscope operating at 200 kV in zero-energy-loss mode with a slit width of 20 eV, and equipped with a 4k × 4k slow-scan CDD camera (Gatan inc.).

## Results

### Diversity of the 22 selected pioneer peptides

To develop our peptide classification method, we took _306_VQIVYK_311_ Tau’s amyloid sequence, also known as ß1 (Fitzpatrick et al., 2017), as a reference. To limit our approach to a feasible number of cases, we chose peptides with the same amino acid composition as the reference β1 (I, K, Q, 2V, Y) but in a different order. To further reduce the targets to a number appropriate for non-automated biochemical analysis, we selected 22 cases (named 2p to 23p, **Table 1**) *via* a combination of approaches and the following criteria: *i*) presence/absence of the hexapeptide in protein sequences referenced in Prosite database, *ii*) presence in different organisms from bacteria to human, and *iii*) predicted amyloid properties. The selected peptides are present in proteins from vertebrates (ß1, 2p, 3p, 4p, 6p, 8p, 9p, 10p, 11p, 12p, 13p, 14p, 17p, 18p, 23p), nematodes (13p, 21p), viruses (3p, 4p, 8p), fungi (6p, 14p, 19p, 21p), a protozoan (23p), a plant (10p), and prokaryotes (4p, 6p, 7p, 8p, 10p, 13p, 15p, 16p, 20p, 21p). One peptide is absent from the Uniprot protein sequence database (22p), while others are contained in a single protein (ß1, 2p, 11p, 12p, 15p, 16p, 18p, 19p) or even up to seven proteins (*e*.*g*. 6p is present in the protein sequences of vertebrates, insects, fungi, and bacteria). Seven peptides are absent from the human proteome (12p, 15p, 16p, 19p, 20p, 21p, 22p). Some are exclusively present in vertebrate proteomes (ß1, 2p, 11p, 12p, 18p) while others are exclusively found in prokaryotic proteomes (5p, 7p, 15p, 16p and 20p).

**Table 1:**
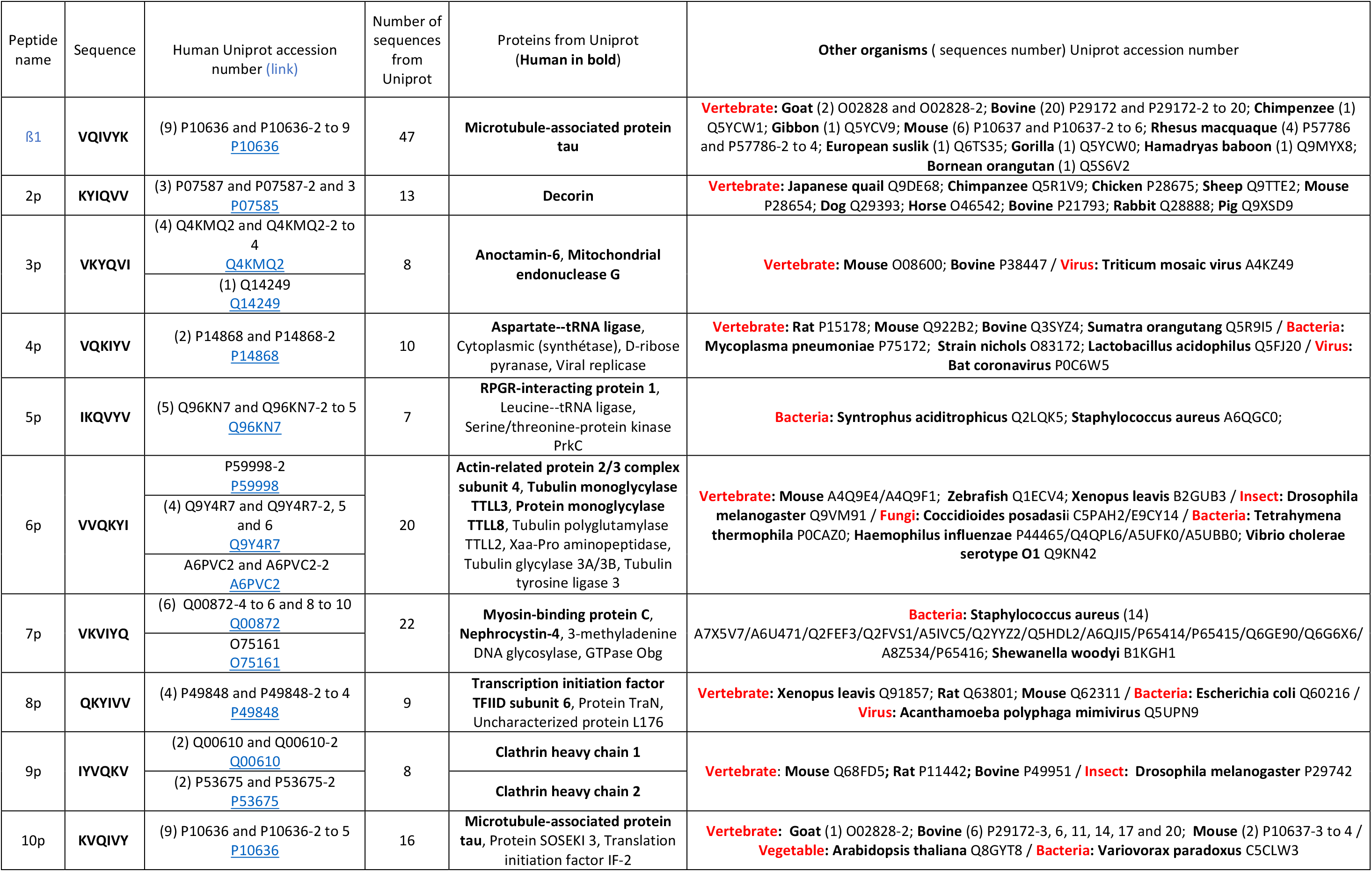
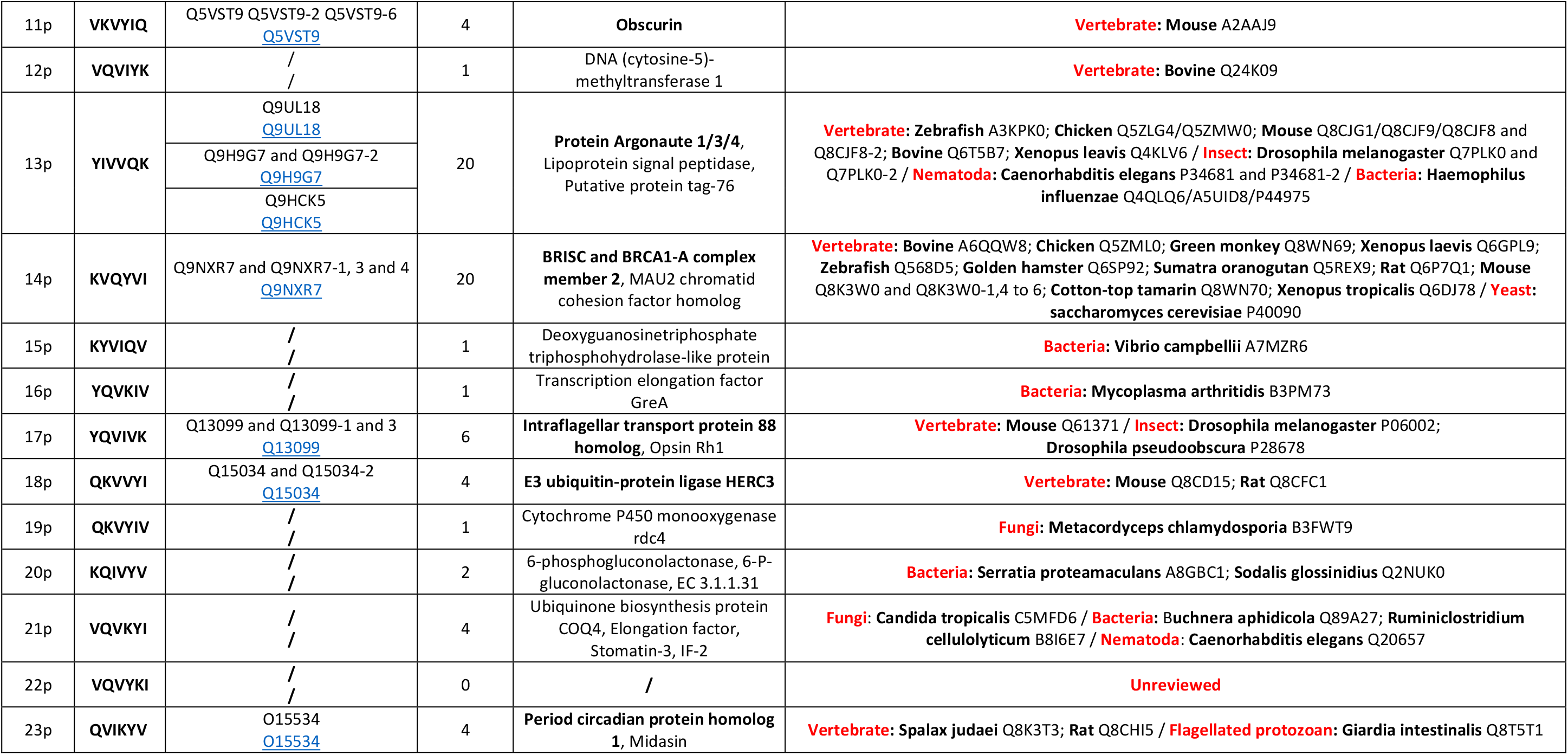
Sequence of hexapeptides studied in this work in relation to their presence in protein sequences in humans and in different organisms.

### Highly heterogeneous predicted amyloidogenicity of similar peptides

We initially employed Tango, Pasta and Salsa to analyze the large virtual peptide comprising target peptides sequences separated by a non-amyloidogenic sequence (see Materials and Methods). We observed a limited variation in amyloidogenicity prediction ranging from 46% to 139% for 6p and 13p, respectively (**Figure 1A**). We found that Salsa was not discriminating enough since it’s amyloidogenic property score varied by approximately 13% between peptides. Thus, we repeated the predictions using only Pasta and Tango that showed the best performance. Prediction of the amyloid character of 22 peptides yielded a robust amplitude varying from 3% to 190% for 3p and 13p, respectively (**Figure 1B**). 13p had a significantly higher score than ß1, 12p and 17p showing an equivalent score and eight peptides (2p, 7p, 8p, 10p,14p, 15p, 18p, 19p and 20p) showing a score 2-4-fold lower than the reference. The remaining peptides showed a low predicted amyloid score (< 25%) or did not show any amyloidogenicity (3p, 4p, 5p, 6p, 21p, 22p, 23p). We conclude that Pasta and Tango yielded homogeneous results, enabling us to classify the peptides according to their amyloidogenic properties.

**Figure 1:**
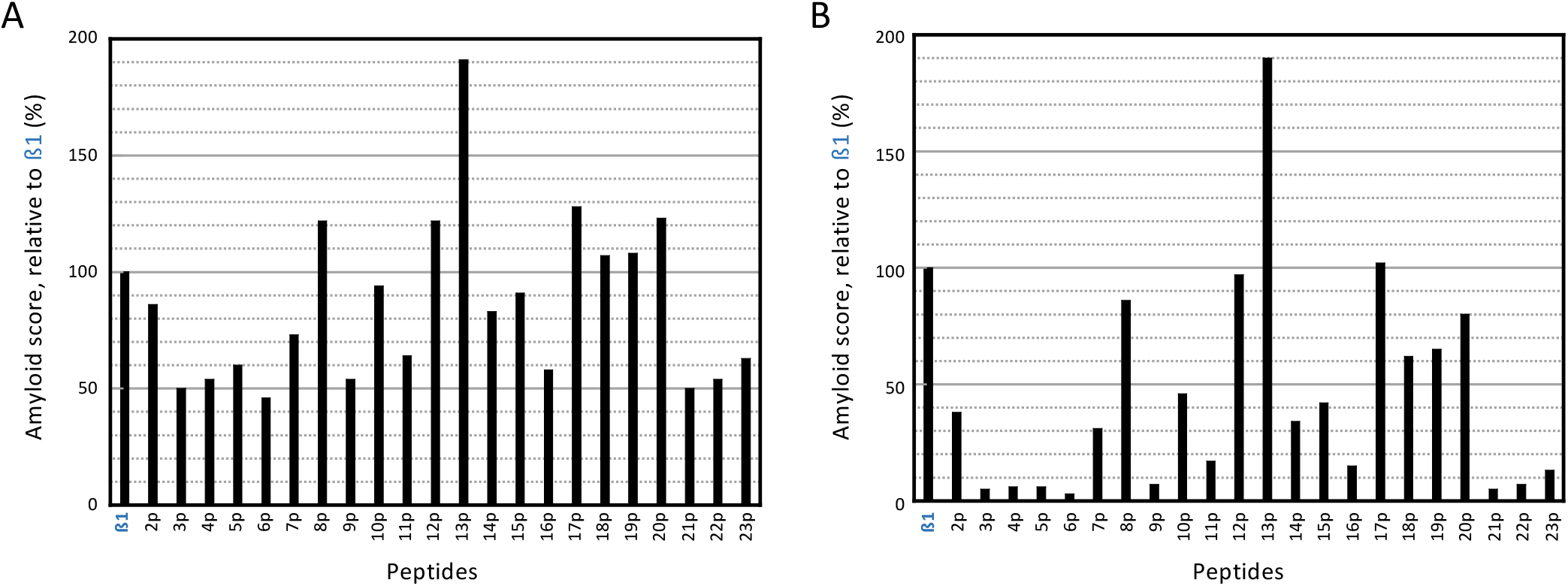
Prediction of the amyloid character of 22 peptides. (A) Results obtained using Tango, Pasta and Salsa prediction methods. (B) Results obtained considering only Pasta and Tango prediction methods. The values were normalized against the reference peptide ß1 VQIVYK (PHF6) for which we set the amyloid score at 100%.

### Target peptide classification

To classify peptides, we quantified their aggregative and amyloidogenic properties and plotted the scores using a graphical representation we named Garnier-Delamarche (GD-plot). The percentage of sedimentable peptide at the working concentration reflects the peptide’s aggregative property, it was reported on the x-axis. The relative fluorescence intensity in the presence of Thioflavin T calculated relative to β1 is a measure for amyloidogenic property, it was reported on the y-axis (**Figure 2**). None of the analyzed peptides exhibited higher amyloidogenicity than the reference peptide ß1. We were unable to study 16p* at 800 μM due to its low solubility in water. Even at a concentration of 288 μM 16p* showed strong aggregation (threshold concentration of ∼10-20 μM); however, it was not amyloidogenic.

**Figure 2:**
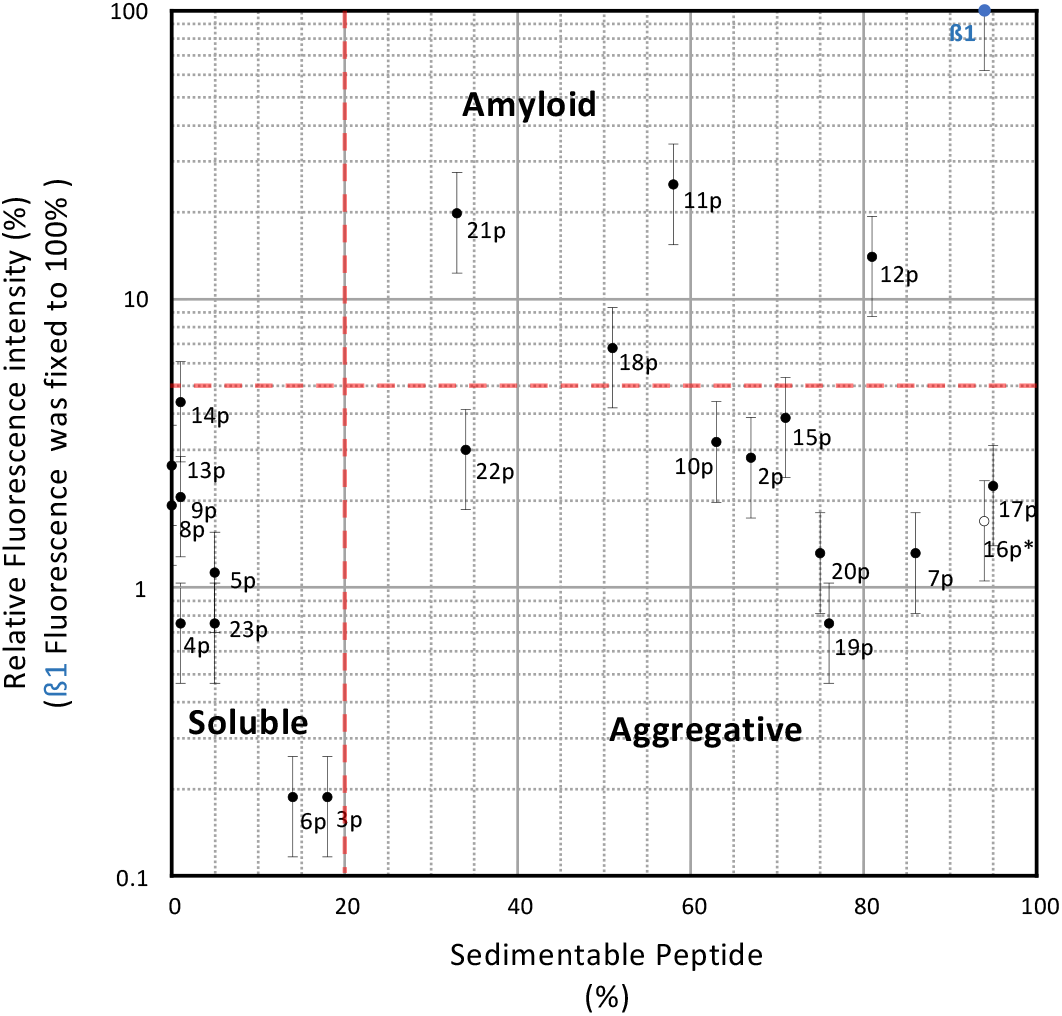
Biochemical analysis and classification of ß1 derivative peptides using a GD-plot. The horizontal and vertical dotted lines correspond to the 5% limit relative to ß1 fluorescence and to the 20% threshold for aggregating peptides, respectively. * 16p was studied at ∼ 290 μM due to its poor solubility in water.

We defined thresholds that depend upon peptides analyzed, the reference peptide and their concentration in the experimental sample. At a concentration of 800 μM, we considered peptides as aggregative beyond 20% of sedimentable structures and as amyloid when the fluorescence exceeded 5% of the fluorescence signal obtained for β1. Hexapeptides 2p, 7p, 10p, 15p, 17p, 19p, 20p, 22p were classified as “aggregative” because of their low fluorescence signal, while 11p, 12p, 18p, 21p were classified as “amyloid”. For instance, 7p and 17p peptides exhibit an aggregative properties equivalent to ß1 (nearly 90%) without fluorescence classifying them as non-amyloidogenic. In the other hand, 11p exhibited ∼60% aggregative properties and showed a ∼ 25% fluorescence intensity compared to ß1, which classifies as amyloidogenic. For peptides showing fluorescence intensity > 5% compared to ß1 (11p, 12p, 18p, 21p), we determined an “amyloidogenicity” factor by calculating the ratio between the fluorescence intensity and the concentration of sedimentable structures (**Table 2**). The standardized factor was determined to be 29, 16, 13, and 38 μM^-1^ for 11p, 12p, 18p, and 21p, respectively. Thus, amyloidogenicity clearly varies between peptides, which underline that Thioflavin T incorporation and/or fluorescence depend on structures formed. Nine peptides were entirely soluble at 800 μM showing very weak (3p, 5p, 6p, 23p) or no (4p, 8p, 9p, 13p, 14p) aggregation properties. We conclude that biochemical results (fluorescence and sedimentation) represented in GD-plot well separate the hexapeptides into three property groups: aggregatives, amyloids and soluble peptides that were neither aggregatives nor amyloids.

**Table 2:**
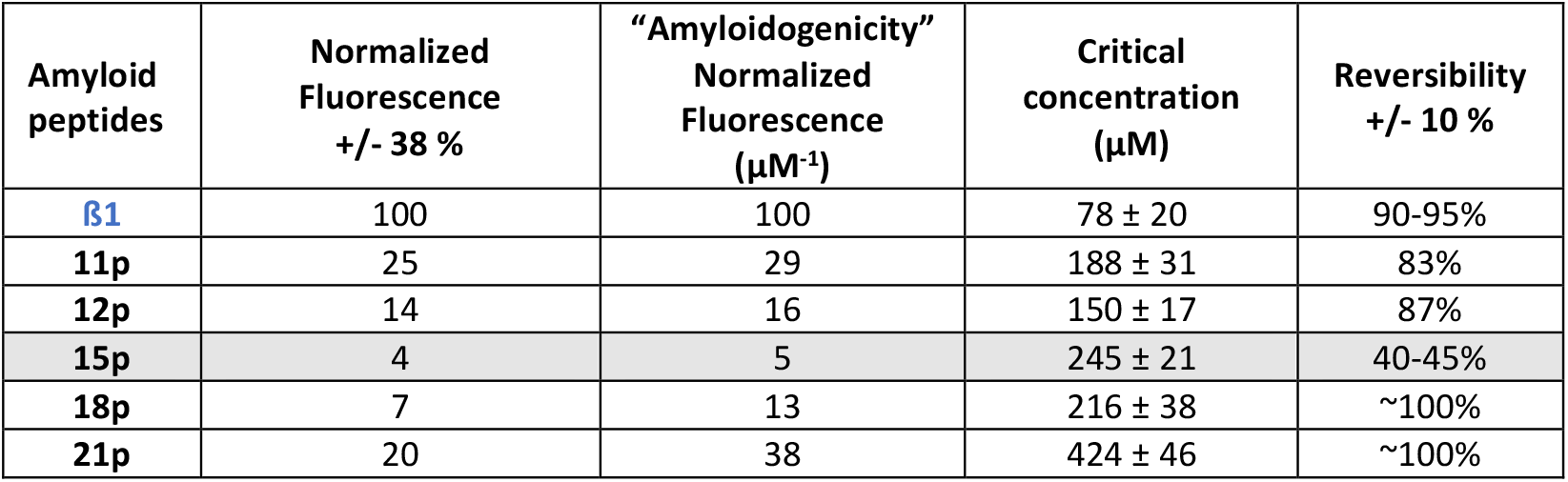
Physical and chemical properties of peptides classified in the amyloid group. The level of “amyloidogenicity” corresponds to the fluorescence intensity relative to the quantity of sedimentable peptide.

### Comparison between biochemical properties and predicted amyloidogenicity

Once each peptide was biochemically characterized, we compared its aggregative and amyloid properties in solution to the predicted *in silico* amyloid predictions. For each peptide, *in silico* prediction was positioned on the abscissa x-axis when compared to its amyloid property (**Figure 3A**), while it was positioned on the ordinates y-axis when compared to its aggregative properties (**Figure 3B**).

**Figure 3:**
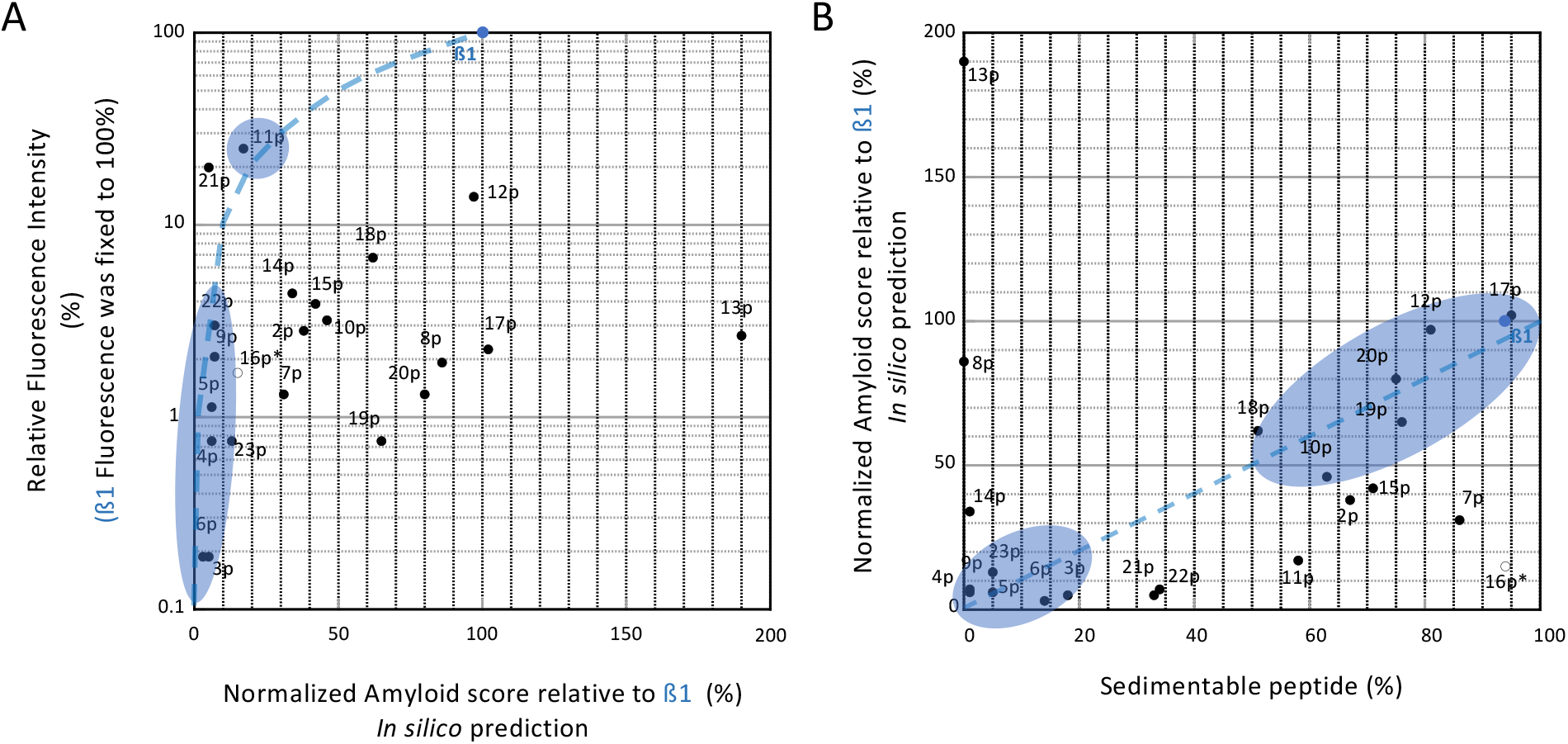
Comparison of biochemical and amyloid properties to in silico prediction of target peptides. **(A)** Plot of amyloidogenic peptide properties determined *in vitro* against the predicted amyloid score (Pasta/Tango) normalized to ß1 is shown. **(B)** Plot of predicted amyloid scores (Pasta/Tango) a plotted against aggregative peptide properties determined *in vitro*. The blue dotted curves correspond to the ideal correlation. Peptides for which the correlation is promising are marked in blue.

The graph in Figure 3A allows for the comparison of peptide amyloidogenicity measured *in vitro* to the predicted amyloid score. Among four peptides showing amyloid properties *in vitro* (11p, 12p, 18p and 21p), only 11p showed a low amyloid feature close to the prediction (∼20-25% compared to ß1). The amyloidogenicity of 12p and 18p was predicted to be close to β1’s, but turned to be much lower and the 21p peptide was predicted to be non-amyloid. In contrast, seven of the peptides predicted to be non-amyloid were soluble *in vitro* (3p, 4p, 5p, 6p, 9p, 22p, and 23p). All other peptides had a predicted relative amyloid score greater than or equal to 30% but they were not found to be amyloid *in vitro*; for example, 19p, 20p, 8p, 17p, whose score was close to ß1, and 13p for which the score was twice as high.

Furthermore, we compared peptides predicted amyloid score to their aggregative properties (**Figure 3B**). The graph highlighted that only for a dozen peptides the predicted amyloid score showed a good correlation with aggregative properties (at ± 10-20%). This was found to be the case for the group of non-aggregative (3p, 4p, 5p, 6p, 9p, 23p), aggregative (10p, 17p, 19p, 20) and amyloid (12p, 18p) peptides. For the remaining peptides, considerable differences between predicted and biochemical properties were observed. For example, 14p, 8p and 13p were predicted to be amyloid but turned out to be entirely soluble. Conversely, predicted non-amyloid peptides showed either a strong aggregation potential (2p, 7p, 15p and 16p) or an amyloidogenic character (11p and 21p). Thus, we observed a poor correlation between the predicted amyloid score and amyloid property determined *in vitro*. This fact notwithstanding, the amyloid score predicted *in silico* was consistent with their sedimentability/aggregative properties measured *in vitro* in about 50% of the targets.

### Assembly properties of amyloid and aggregative peptides: critical concentration, aggregation threshold and reversibility

Since we are interested in amyloid peptides, we studied the β1, 11p, 12p, 18p and 21p of the amyloid group and 15p which showed 70% aggregation and a fluorescence signal of approximately 4% compared to β1 (**Figure 2**). 15p is interesting because its fluorescence signal was close to the 5% threshold distinguishing aggregative from amyloids peptides. First, we determined the Cr of these 6 peptides by centrifugation (**Figure 4** and **Table 2**). As expected, we observed a negative correlation between the quantity of sedimented structures at 800μM (**Figure 2**, x-axis) and the Cr. This was true except for the 15p as compared to 11p and 18p peptides. Indeed, 15p, which seems to be more aggregative had a higher Cr than 11p and 18p. This could be explained by the difference in slope between Cr linear regressions. Indeed, 15p showed a slope of 1.1 whereas the slopes were of 0.74 and 0.84 for 11p and 18p, respectively. This explained why at high concentrations (**Figure 2**), the quantity of sedimentable structures tended to be lower for 11p and 18p than for 15p. For peptides classified as non-amyloid (aggregative or soluble), we did not determine a critical concentration but an aggregation threshold (**Table 3**).

**Table 3:** Peptide solubility/aggregation thresholds. The peptides considered to be soluble (reported in Larralde et al., n.d.) are highlighted in grey. * threshold calculated considering the studied concentration of 290 μM.

| Peptides | Aggregation Threshold<br>($\mu\text{M}$) | Peptides | Aggregation Threshold<br>($\mu\text{M}$) |
| --- | --- | --- | --- |
| 2p | 264 | 13p | > 800 |
| 3p | 656 | 14p | > 800 |
| 4p | 792 | 15p | 220 |
| 5p | 760 | 16p* | 17 |
| 6p | 688 | 17p | 40 |
| 7p | 112 | 19p | 192 |
| 8p | > 800 | 20p | 200 |
| 9p | > 800 | 22p | 528 |
| 10p | 296 | 23p | 760 |

**Figure 4:**
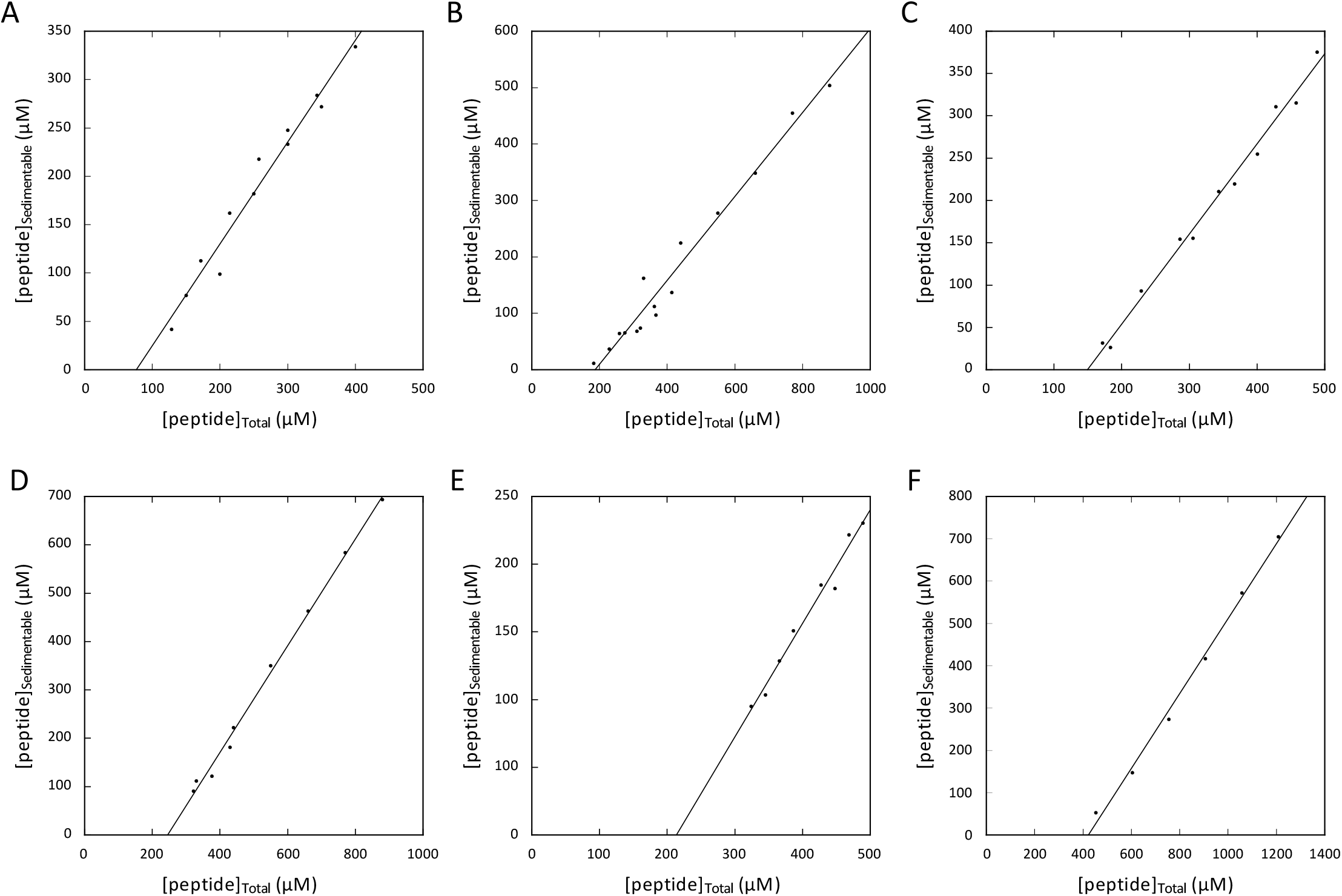
Critical assembly concentration (Cr) of amyloid peptides as determined by centrifugation. (A-F) Graphs show CRs of ß1, 11p, 12p, 15p, 18p, and 21p. Total versus sedimented peptide concentrations in micromoles (μM) are plotted on the x- and y-axes, respectively.

To further characterize the structures formed by our target peptides, we developed a protocol to study their reversibility by dilution. Indeed, the speed and percentage of reversion varied depending on the polymerization concentration, the final concentration after dilution, and the reversibility of the associative process. Peptide polymerization was induced at a concentration equal to twice the Cr and then samples were diluted 8-fold to reach a final concentration of 0.25 Cr (see Materials and Methods, **Figure 5** and **Table 2**). As could be seen in the case of ß1, diluting the sample was accompanied by a strong decrease of Thioflavin T fluorescence intensity, reflecting the loss of cross-beta structures and therefore the reversion of the amyloid structures. The reversion process reached 90-95% after 40 minutes (**Figure 5**). An almost total reversion was observed for the other peptides classified as amyloids (11p, 12p, 18p and 21p; **Table 2**).

**Figure 5:**
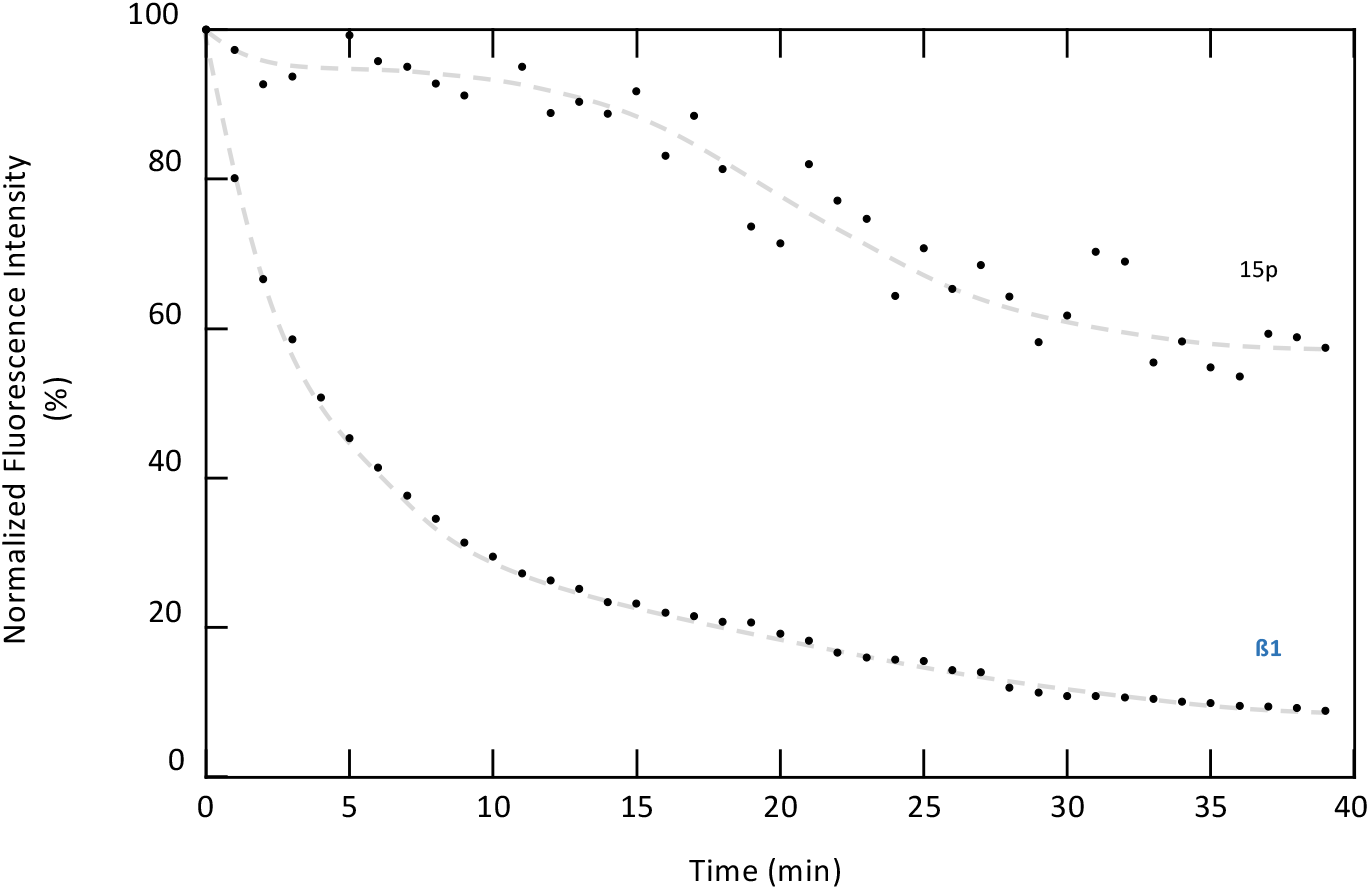
Reversibility of amyloid peptides structure. A graph shows fluorescence intensities (y-axis) plotted against time (x-axis) for target and reference peptides as indicated. The gray dotted curves show the progression of the depolymerization process.

Contrary to amyloid peptides, depolymerization of 15p showed a slight decrease in Thioflavin T fluorescence intensity after dilution, which had a sigmoid appearance. The decrease in fluorescent intensity was almost zero during the first twenty minutes, then accelerated to tend towards a plateau not exceeding 40 to 50% reversion after 40 minutes (**Figure 5** and **Table 2**). The fluorescence threshold arbitrarily set at 5% fluorescence (**Figure 2**, GD-plot) to distinguish amyloid from aggregative peptides reveals its efficiency here. Indeed, the structures formed by 15p (normalized fluorescence 5 μM^-1^, **Table 2**) behaved more like irreversible aggregates than like reversible amyloid structures.

### Heterogeneity of structures formed by peptides with auto associative properties

To identify the characteristic signatures for the different self-associative peptides (aggregative *versus* amyloid), we analyzed structures by negative staining transmission electron microscopy. In addition to self-associative peptides, we also studied the soluble peptide 9p. (**Figure 6, Table 4**). Structures ranging from crystalline, to more or less compact flat or twisted ribbons/fibers were observed. β1 formed characteristic homogeneous compact fibrillar structures with a diameter of ∼10 nm and a pitch of ∼100 nm. ß1, 7p, 11p, 12p, 18p, 19p, 20p and 22p also formed fibrillar structures with regular or irregular diameters ranging from 5 to 20 nm and a twist pitch of 100 to 200 nm (**Figure 6** and **Table 4**). 2p, 10p, 15p, 16p and 17p showed structures resembling ribbons with widths of around twenty nanometers. 21p formed globular/ovoid structures. Furthermore, the 9p soluble peptide showed crystal structures, the images obtained by Cryo-EM confirmed the absence of structures (data not shown).

**Table 4:**
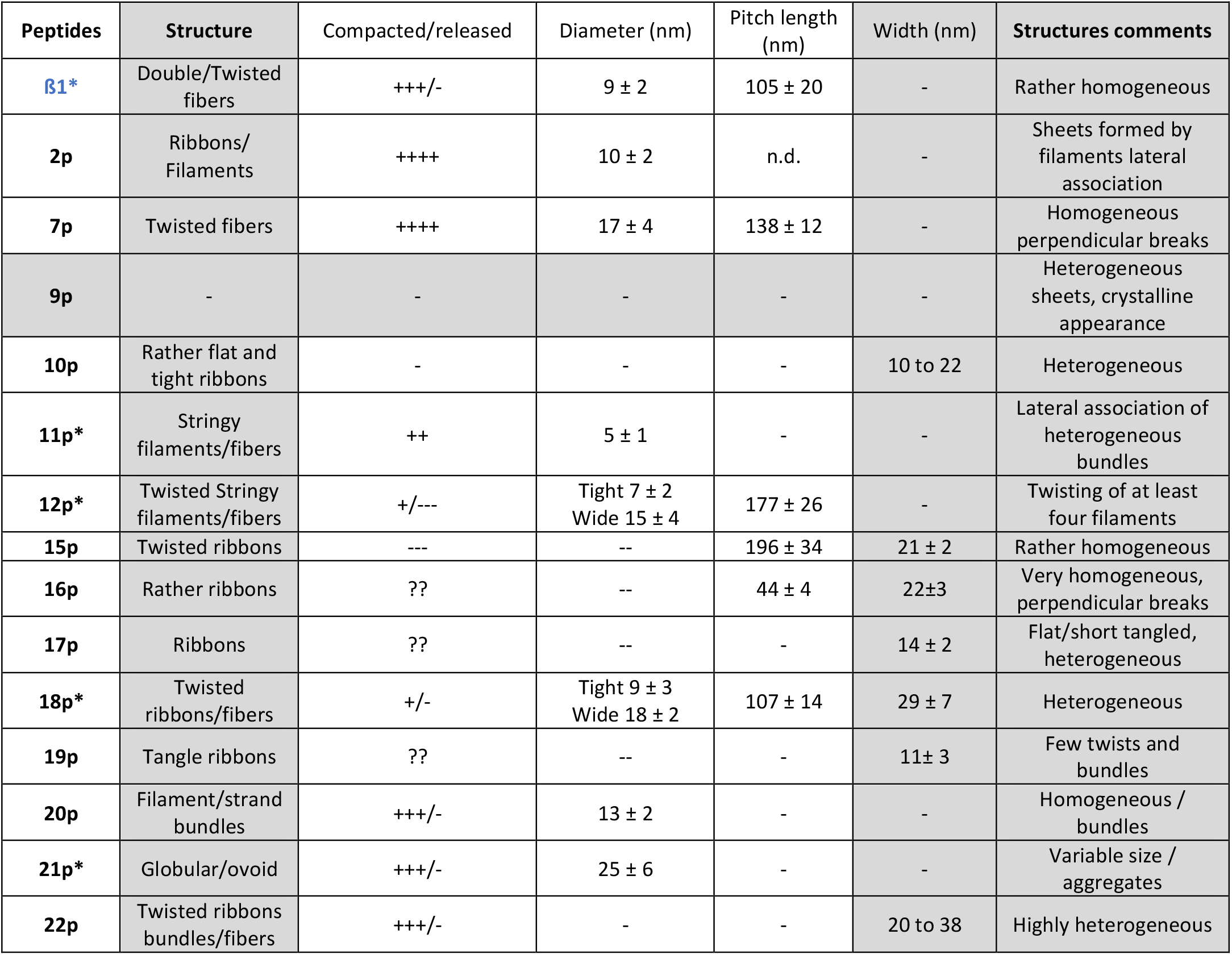
Analysis and comments on the structures formed by peptides belonging to the amyloid and aggregative groups.

**Figure 6:**
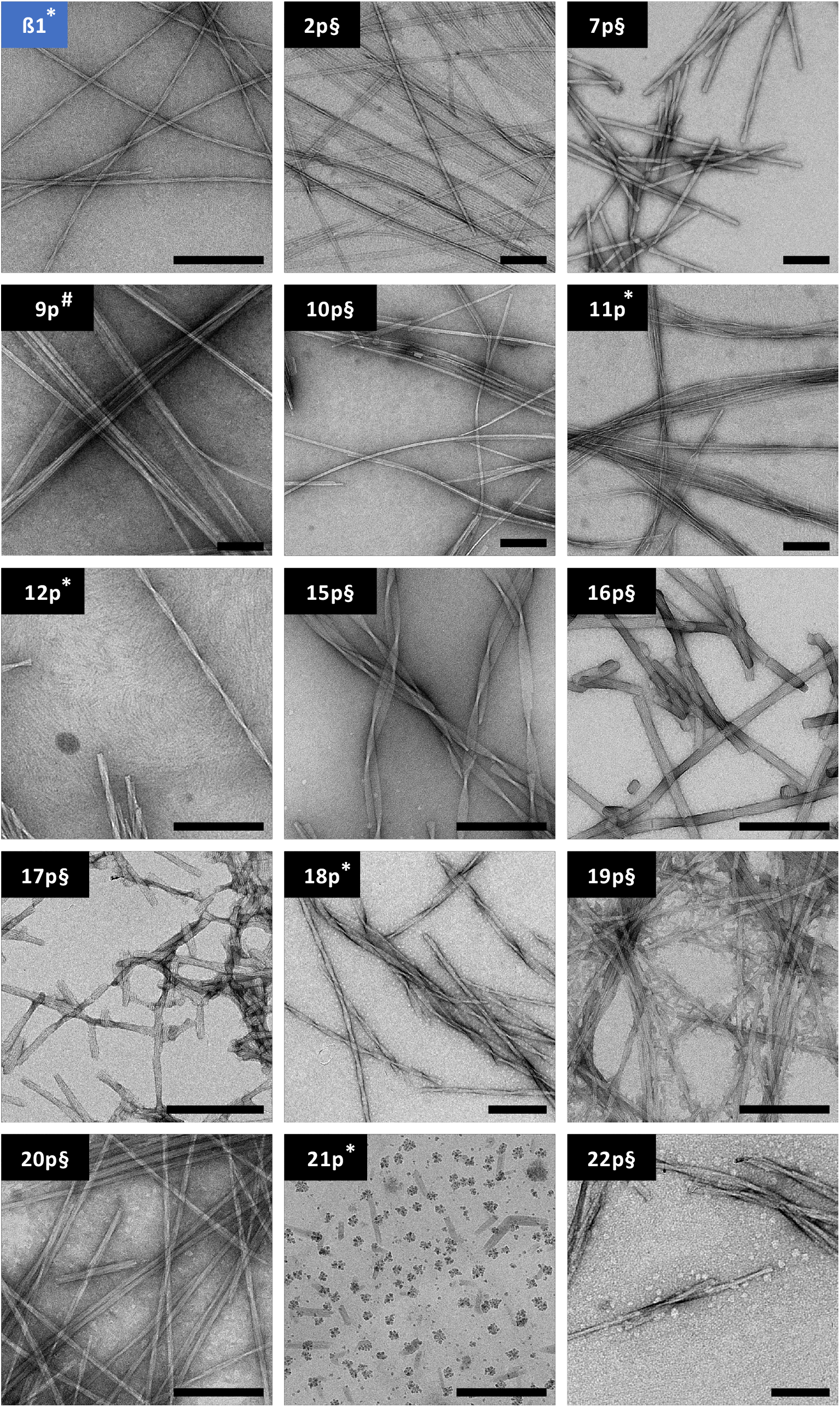
Negative staining TEM analysis of structures formed by peptides. *Peptides belonging to the amyloid (*), aggregative (§) and soluble (#) groups*. The characteristics of the structures were summarized in Table 4. Scale bar 200 nm.

We conclude that while amyloid and aggregative peptides can form fibers. Apart from 21p, all selected hexapeptides followed a similar auto-associative process corresponding to a primary polymerization in the form of a ribbon, which, depending on the imperfections and the steric hindrance of the different amino acids, tends to twist forming more or less compact filaments or fibers. We were unable to identify any structure showing an amyloid signature via TEM imaging.

## Discussion

Amyloidosis are pathologies resulting from the disruption of the cellular or extracellular homeostasis of one or more proteins which, following their loss of function, acquire new capacities to form amyloid fibers. Most often, these are self-associative processes involving only part of the protein sequence (a few dozen amino acids) which, ultimately, end up compacted in the core of the fibers (Scheres et al., 2023). Within these protein sequences are found short peptides (not exceeding 5 to 10 amino acids) responsible for the nucleation processes at the origin of fiber formation (Sgourakis et al., 2007; Smit et al., 2017). These short amyloid peptides, being (or becoming) accessible in the structure of the original protein, self-associate in the form of beta sheets, often parallel, allowing to the establishment of intermolecular bonds between proteins. Once these initial sheets are formed, they are capable of establishing long-distance intramolecular interactions with other sequences of the protein by interdigitation of the side chains. A double zipper-like propagation occurring at both intermolecular beta sheets and intramolecular interdigitation of the side chains leads to protein on itself folding perpendicular to the axis of the fiber. The identification and classification of these short nucleator peptides and related peptides are therefore of primary interest for understanding these nucleation phenomena developing therapeutic strategies.

To tackle this critical problem, we developed a simple biochemical method for the analysis of a short amyloid reference peptide and its related peptides with the aim of predicting aggregative/amyloid properties used to classify them. For each peptide we determined their aggregative properties in a physiological environment (reflecting their solubility), and their capacity to initiate a fluorescence signal in the presence of Thioflavin T (reflecting their amyloid features). These parameters position peptides in a GD-plot to classify them.

To apply and validate our method of analysis and classification, we chose ß1 (PHF6, VQIVYK), named ß1 in Tau protein’s fibers (Fitzpatrick et al., 2017), as a reference amyloid peptide (Perez et al., 2007; Smit et al., 2017). The biochemical analysis of our batch of peptides required determining a working concentration for all peptides, which depends on the properties of the reference peptide and the solubility of the target peptides when dissolved in water. Since all peptides, except 16p, showed a water solubility threshold between 1 and 1.5 mM, we chose the concentration of 800 μM. This was close to their solubility threshold while maintaining a safety margin. After measuring the biochemical parameters, the peptides were analyzed using GD-plots.

Among the peptides studied, none showed an amyloidogenic character greater than ß1. The analysis of another batch of peptides revealed that some could show fluorescence intensities 3-4 fold higher than ß1 (Larralde et al., n.d.). Thus, because of the large disparity in fluorescence observed between the studied peptides (< 1% here and > 300% in (Larralde et al., n.d.)), we chose to display data on a logarithmic scale on the ordinate to avoid overwriting the points on the x-axis of the GD-plot. Using this graphical representation, we identified three peptide groups: purely aggregative, amyloid and soluble peptides. We arbitrarily set thresholds for solubility at 20% and fluorescence at 5% in the context of the peptide’s working concentration and peptide batch analyzed. As far as solubility is concerned, the threshold is not crucial since we know the solubility limit for each peptide. By setting it at 20%, all peptides in this group are soluble up to a concentration of 640 μM. For amyloidogenicity, the 5% threshold was more difficult to determine. We demonstrated that all peptides classified as amyloid exhibited complete reversibility of structures formed whereas for 15p, classified as aggregative, the reversibility was only partial. Thus, depending on the batch of peptides studied, reversibility criteria could be used to set a threshold making it possible to discriminate amyloid from aggregative peptides. To compare the fluorescence of amyloid peptides, we determined a normalized amyloidogenicity parameter, which is expressed in μM^-1^ and corresponds to the relative fluorescence intensity related to the concentration of sedimentable structures. The normalized amyloidogenicity clearly shows that between the different peptides, the fluorescence intensity of Thioflavin T is not proportional to the quantity of sedimentable structures. This demonstrates that Thioflavin T does not intercalate into different amyloid with the same efficiency, which explains varying fluorescence signal intensities. Here, all the peptides present a weaker amyloid feature than ß1. However, we observed the opposite effect in work on a different batch of peptides (Larralde et al., n.d.). TEM analysis did not reveal characteristic structures of amyloid peptide because the morphologies range from flat/twisted ribbons to relaxed/compact fibers were observed for both amyloid and aggregative peptides.

## Conclusion

The approach to peptide classification we describe here is a simple and rapid biochemical analysis method. Our method can be automated by optimally coupling measurements of sample fluorescence with the quantification of sedimentable structures. Peptide classification can be used to design other batches of peptides related to a reference peptide by modifying/conserving the position of certain key amino acids. While aggregative peptides are of limited interest, it is not the case for amyloid and soluble peptides. Indeed, the highlighting of new amyloid peptides in protein sequences allows to identify new nucleation centers responsible of pathological fibers and a better understanding of their amyloid self-association process (kinetics and structure determination) could lead to develop new therapies. Likewise, soluble peptides are of clinical interest because they are potential inhibitors of the nucleation process of the reference peptide responsible for the formation of pathological fibers and because they are quite easy to administer (Larralde et al., n.d.).

## Acknowledgments

We thank Mike Priming for critical reading of the manuscript and for his scientific expertise.

